# Targeting MEK5 impairs non-homologous end-joining repair and sensitizes prostate cancer to DNA damaging agents

**DOI:** 10.1101/737080

**Authors:** Constantinos G. Broustas, Axel J. Duval, Kunal R. Chaudhary, Richard A. Friedman, Renu K. Virk, Howard B. Lieberman

## Abstract

Radiotherapy is commonly used to treat a variety of solid human tumors, including localized prostate cancer. However, treatment failure often ensues due to tumor intrinsic or acquired radioresistance. Here we find that the MEK5/ERK5 signaling pathway is associated with resistance to genotoxic stress in aggressive prostate cancer cells. MEK5 knockdown by RNA interference sensitizes prostate cancer cells to ionizing radiation (IR) and etoposide treatment, as assessed by clonogenic survival and short-term proliferation assays. Mechanistically, MEK5 downregulation impairs phosphorylation of the catalytic subunit of DNA-PK at serine 2056 in response to IR or etoposide treatment. Although MEK5 knockdown does not influence the initial appearance of radiation- and etoposide-induced γH2AX and 53BP1 foci, it markedly delays their resolution, indicating a DNA repair defect. A cell-based assay shows that non-homologous end joining (NHEJ) is compromised in cells with ablated MEK5 protein expression. Finally, MEK5 silencing combined with focal irradiation causes strong inhibition of tumor growth in mouse xenografts, compared with MEK5 depletion or radiation alone. These findings reveal a convergence between MEK5 signaling and DNA repair by NHEJ in conferring resistance to genotoxic stress in advanced prostate cancer and suggest targeting MEK5 as an effective therapeutic intervention in the management of this disease.

## Introduction

Radiotherapy is a common therapeutic modality for the treatment of human epithelial tumors, including those of prostate origin [1]. Despite considerable improvements in delivering the radiation dose with precision, therapeutic benefit in prostate cancer radiotherapy has been hampered by tumor resistance to ionizing radiation. Tumor-intrinsic pro-survival pathways, as well as upregulation of DNA repair pathways constitute major mechanisms by which malignant cells become radioresistant [2].

Cells react to genotoxic insults by engaging a highly intricate DNA damage response and repair network, which is mediated by the phosphoinositide-3-kinase-like kinases (PIKKs) DNA-PK (DNA-dependent protein kinase), ATM (ataxia telangiectasia mutated), and ATR (ATM and Rad3-related) [3]. DNA-PK and ATM are activated by DSBs, whereas ATR plays a leading role in response to DNA single-strand breaks [3]. DNA double strand breaks (DSBs) induced by ionizing radiation or certain chemotherapeutic agents potentially represent a highly toxic form of DNA damage that leads to cell death or genomic instability. In mammals, there are two major pathways for repairing DSBs. Homologous recombination (HR) is predominantly error-free repair and active during the S and G2 phases of the cell cycle, and non-homologous end-joining (NHEJ) that can be either error-free or error-prone and is active throughout the cell cycle [4, 5]. NHEJ is the dominant pathway for repairing DNA DSBs in mammalian somatic cells [6]. Central to NHEJ repair is the DNA-PK trimeric complex, composed of DNA-PK catalytic subunit (DNA-PKcs) and DNA binding subunits, KU70 and KU80. Both KU70 and KU80 bind to DNA breaks and activate DNA-PKcs kinase activity to initiate DNA repair by NHEJ [7]. Phosphorylation at Threonine 2609 (S2609) and Serine 2056 (S2056) in response to DNA DSBs is associated with repair efficiency of DNA-PKcs [8].

Mitogen-activated protein kinase kinase 5 (MAP2K5 or MEK5) belongs to the family of MAP kinases. It is activated by the upstream kinases MEKK2 and MEKK3 at serine 311 and threonine 315 (S311/T315), or in some cases directly by c-Src [9–12]. MEK5, in turn, phosphorylates and activates extracellular signal-regulated kinase 5 (ERK5 or BMK1) at T218/Y220 [9]. The MEK5/ERK5 pathway can be activated by various stimuli such as oxidative stress, growth factors, and mitogens downstream of receptor tyrosine kinases, as well as G protein-coupled receptors, and culminates in the activation of a large number of transcription factors, including MEF2 (myocyte enhancer factor 2), c-JUN, NF-κB, and transcription factors that control the epithelial-mesenchymal transition (EMT) program [13–18]. Furthermore, recent reports have shown that ERK5 is activated by oncogenic BRAF and promotes melanoma growth [19], whereas inhibition of ERK1/2 in melanoma leads to compensatory activation of the MEK5/ERK5 pathway [20].

The MEK5/ERK5 pathway plays a pivotal role in prostate cancer initiation and progression. MEK5 protein is overexpressed in prostate cancer cells compared with normal cells and MEK5 levels are correlated with prostate cancer metastasis [21]. Furthermore, high expression of ERK5 in prostate cancer has also been found to correlate with poor disease-specific survival and could serve as an independent prognostic factor [22]. Moreover, ERK5 expression in prostate cancer is associated with an invasive phenotype [23]. Recently, it has been shown that deletion of *Erk5* in an established *Pten*-deficient mouse model of human prostate cancer can increase T-cell infiltration and control tumor growth [24].

The present study was designed to investigate whether MEK5 downregulation sensitizes human prostate cancer cells to radiation and other agents that inflict DNA DSBs, and examine the potential mechanism of sensitization to these drugs. We show that MEK5 knockdown enhances the sensitivity of human prostate cancer cells to radiation and etoposide, which, mechanistically, can be attributed to inhibition of DNA-PKcs phosphorylation and the non-homologous end-joining process. Importantly, *in vivo* studies using a mouse xenograft model show that MEK5 ablation synergizes with radiation to suppress tumor growth. Our results support the hypothesis that inactivation of MEK5 in prostate cancer could be a strategy for improving the efficacy of radiotherapy in prostate cancer patients.

## Results

### MEK5/ERK5 pathway activation in response to ionizing radiation

It has been demonstrated previously that MEK5 and ERK5 are upregulated in human prostate cancer and are associated with metastasis and reduced patient survival [25–27]. Immunoblotting of a panel of normal and malignant human prostate cell lines showed that MEK5 is predominantly expressed in advanced prostate cancer cell lines PC3 and DU145, less in androgen-responsive LNCaP, and at very low levels in normal epithelial prostate cells (PrEC) and the immortalized, but non-tumorigenic, cell line EP156T (Supplemental Fig. 1).

The MEK5/ERK5 pathway is activated by a diverse array of growth factor, cytokines, as well as stress in the form of osmotic stress. We sought to determine whether the MEK5/ERK5 pathway is activated in response to ionizing radiation (IR) in human prostate cancer. Using phospho-ERK5 (T218/Y220) levels as a readout for the activation of the pathway, we exposed DU145 expressing either *MEK5* or control *Luciferase* siRNA to different doses of IR and lysed the cells 15 min post-irradiation. As shown in Fig. 1a, phospho-ERK5 levels were increased after 2 and 4 Gy of γ-rays. We repeated the experiment by exposing PC3 cells to 3 Gy of IR and lysing the cells at various times post-irradiation. Activation of ERK5 in response to IR was fast occurring already at the earliest examined time (5 min) and persisting up to 15-30 min, gradually diminishing at later time points (Fig. 1b). As expected no phospho-ERK5 was detected in the MEK5 depleted DU145 of PC3 cells. Similarly, PC3 cells stably expressing *MEK5* shRNA had reduced levels of activated ERK5, while control cells, stably expressing a scrambled shRNA, showed increase in phospho-ERK5 (T218/Y220) at 10 min post-irradiation that returned to basal levels by 4 h (Fig. 1c). We conclude that ionizing radiation induces a fast and transient activation of MEK5/ERK5 signaling pathway.

**Fig. 1.**
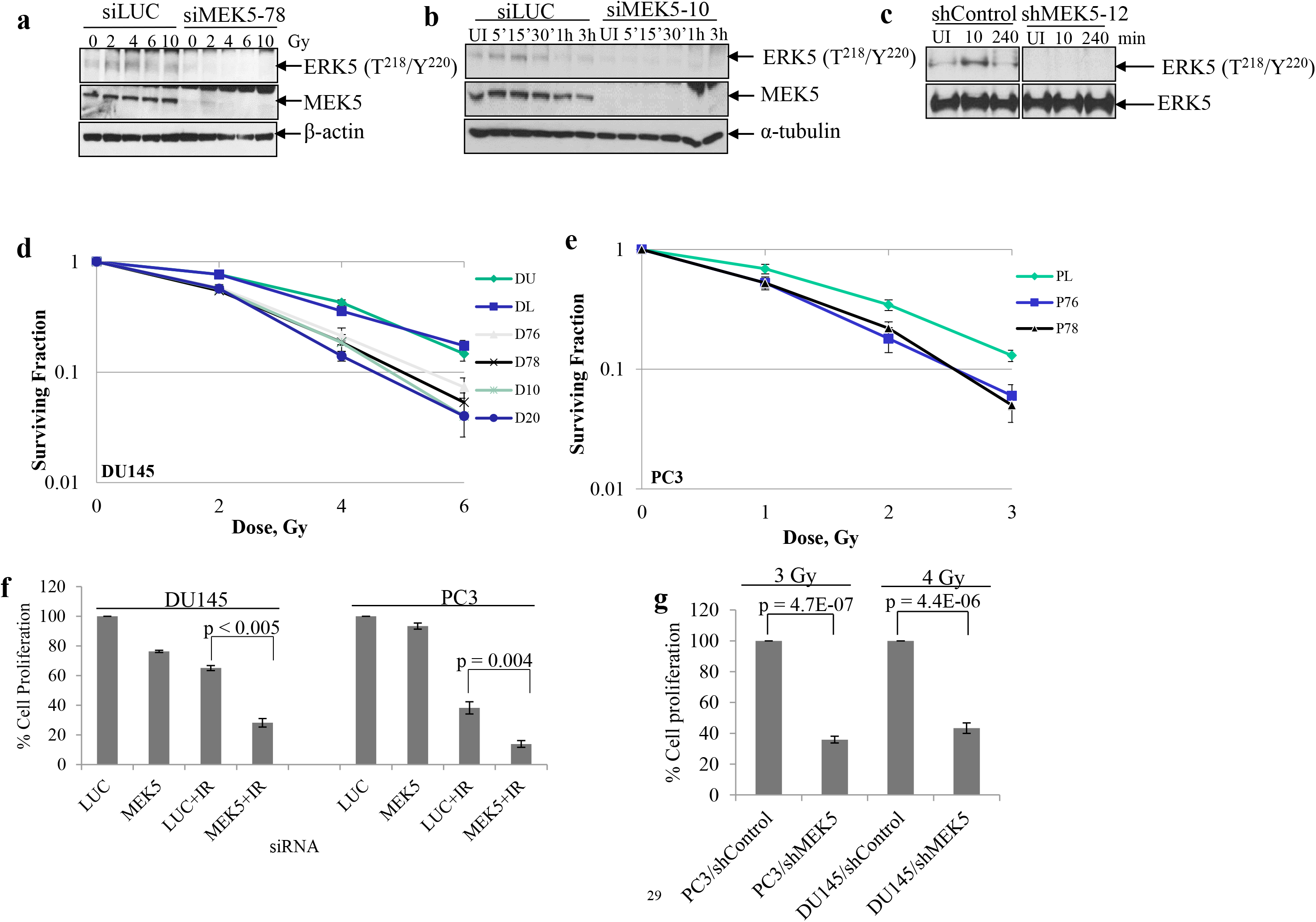
MEK5 silencing sensitizes cells to radiation. **a** DU145 cells were transiently transfected with *Luciferase* (siLUC) or *MEK5* (siMEK5-78) siRNA. Two days later, cells were serum-starved for 24h and irradiated by various doses of γ-radiation. Fifteen minutes later, cells were lysed and proteins were subjected to immunoblotting with the indicated antibodies. **b** Time course activation of ERK5 in response to ionizing radiation. PC3 cells were transiently transfected with either *Luciferase* (siLUC) or *MEK5* (siMEK5-10) siRNA and serum starved for 48h. Cells were irradiated with 4 Gy and lysed at the indicated time points. Levels of total MEK5 and α-tubulin are shown. **c** PC3 stably expressing a scrambled (shControl) or *MEK5* (clone#12) shRNA were irradiated with 3 Gy γ-rays and immunoblotted subsequently with phospho-ERK5 and total ERK5 antibodies. **d** DU145 clonogenic survival assay. DU145 cells were either left untransfected (DU) or transiently transfected with luciferase siRNA (DL) or four different siRNAs against MEK5 (D76, D78, D10, D20). Two days later, cells were irradiated with increasing doses of γ-radiation and plated for clonogenic assay. After 11 days, colonies were scored and normalized against plating efficiency. **e** PC3 cells were transfected with luciferase siRNA (PL) as control or MEK5 siRNAs (P76, P78) and clonogenic assay was carried out as in d. **f** Cell proliferation assay. DU145 and PC3 cells were transiently transfected with control *Luciferase* (LUC) or *MEK5* siRNA. Three days later, cells were irradiated with 4 Gy γ-rays and incubated for 6 days. Cells were trypsinized and counted with a hemocytometer. **g** DU145 and PC3 cells were stably expressing either scrambled (shControl) or *MEK5* (shMEK5) shRNA were treated as in f. Data for d, e, and f represent the mean ± S.D (n = 3). P-values were calculated by Student’s t-test. UI: unirradiated.

### Clonogenic survival assay

To assess the physiological significance of IR-induced MEK5/ERK5 pathway activation, we next assessed the ability of MEK5 depletion to radiosensitize human prostate cancer cells using clonogenic survival assays. For this purpose, we transiently depleted MEK5 from DU145 (four non-overlapping siRNA against MEK5) or PC3 (two independent siMEK5) and two days later irradiated cells with a range of γ-rays. siRNA treatment was able to suppress MEK5 protein levels for at least 7 days (Supplemental Fig. 2a, b). The number of radioresistant clones was recorded in control cells (transfected with *Luciferase* siRNA) and compared with MEK5-depleted cells. *MEK5* knockdown led to significant reproductive cell death after irradiation compared with irradiated cells transfected with *Luciferase* siRNA (Fig. 1d, e). Specifically, knocking down MEK5 by each of four non-overlapping siRNAs sensitized DU145 cells to radiation (surviving fraction at 2 Gy [SF_2_] 0.54 ± 0.02) compared to either parental cells or cells transfected with control luciferase siRNA (SF_2_ 0.78 ± 0.05) (Fig. 1d). Similar radiosensitization was achieved with PC3 cells (SF_2_ 0.35 ± 0.04 vs. 0.20 ± 0.03 in control vs. siMEK5) (Fig. 1e).

We also performed shorter-term cell proliferation assays with PC3 and DU145 cells transiently expressing *MEK5* or *Luciferase* siRNA irradiated or not with 4 Gy, and cells were counted 6 days later. Transfection of untreated PC3 or DU145 cells with siMEK5 did not affect cell proliferation, appreciably. However, cells with MEK5 knockdown showed marked radiosensitization. Thus, cell proliferation of irradiated DU145 cells expressing control siRNA was reduced to 65.1 ± 1.7% (n = 3), whereas in MEK5 knockdown DU145 cells proliferation was 28.2 ± 2.9% (n = 3; p < 0.005). Likewise, irradiated PC3 cells expressing *Luciferase* or *MEK5* siRNA showed 38.3 ± 4.1% (n = 3) and 13.9 ± 2.3% (n = 3) (p < 0.004), respectively (Fig. 1f). Next, we established PC3 and DU145 cells stably expressing *MEK5* or scrambled (shControl) shRNA and isolated 2 clones (#12, #22) for PC3 and 3 clones (denoted #5, #7, and #9) for DU145 cells that showed downregulation of endogenous MEK5 protein (Supplemental Fig. 2c, d). We exposed PC3/shMEK5#12 and DU145/shMEK5#9 cells to 3 Gy (PC3) or 4 Gy (DU145) of γ-rays. In agreement with the clonogenic assay results, silencing of *MEK5* resulted in significant radiosensitization in both PC3 (35.9 ± 2.2%; n = 3; p = 4.7E-07) and DU145 (43.4 ± 3.4%; n = 3; p = 4.4E-06) cells 6 days post-irradiation compared with shControl cells (Fig. 1g and Supplemental Fig. 3 for an additional independent experiment).

### Cell cycle checkpoint activation in response to IR is not affected by MEK5

Cells exposed to genotoxic stress, such as IR, arrest the cell cycle at various phases and attempt to repair the DNA damage. In particular, cells that lack a functional p53, such as PC3 and DU145, arrest the cell cycle at G2/M phase. To determine the impact of MEK5 knockdown on cell cycle checkpoint activation after irradiation we analyzed cell cycle distribution. As expected, irradiation of either DU145 or PC3 cells caused a G2/M arrest starting at about 8 h post-IR, whereas by 48 h cells had resumed their normal cell cycle activity. However, transient or stable downregulation of MEK5 did not appreciably affect cell cycle distribution after irradiation (Supplemental Fig. 4a, b). These results suggest that MEK5 does not play a role in enforcing cell cycle checkpoint activation in response to IR.

### DNA-PKcs activation in response to genotoxic stress is compromised in MEK5 knockdown cells

Many studies have linked defects in DNA repair mechanisms to enhanced radiosensitivity. DNA double strand breaks inflicted by ionizing radiation, etoposide, and other anticancer agents lead to activation of kinases ATM and DNA-PKcs that initiate DNA repair. Activation of ATM is primarily monitored by phosphorylation of serine 1981 (S1981) and ATM is pivotal in the activation of DNA repair by homologous recombination. DNA-PKcs contains multiple Ser/Thr phosphorylation sites, and its DNA damage-inducible autophosphorylation site at S2056 is required for the repair of double-strand breaks by NHEJ [8]. Phosphorylation of serine 2056 (S2056), along with phosphorylation at threonine 2609 (T2609), are considered markers for DNA-PKcs activation in response to DNA damage [7]. Thus, to investigate the potential molecular mechanisms underlying the enhanced sensitivity of MEK5 knockdown in prostate cancer cells to IR, we examined phosphorylation status of DNA-PKcs and ATM in response to DNA damage. PC3 cells transiently expressing a control *Luciferase* siRNA (siLUC) responded to 3 Gy γ-rays by a robust increase in phosphorylation of DNA-PKcs at S2056 and T2609 that was detectable at the earliest time point examined (15 min) post-irradiation. DNA-PKcs phosphorylation signal was diminished to near basal levels 3 h post-irradiation, suggesting completion of DNA repair [7]. In contrast, DNA-PKcs phosphorylation was severely diminished in MEK5 depleted PC3 cells (Fig. 2a). On the other hand, ATM phosphorylation at S1981 in response to IR was comparable between control and MEK5-depleted PC3 cells (Fig. 2a). These results were also confirmed by using DU145 cells (Fig. 2b). We also examined phosphorylation of DNA-PKcs at S2056 using PC3 cells stably expressing MEK5 shRNA (clones #12, #22) or control shRNA (shControl) (Fig. 2c and Supplemental Fig. 5), as well as DU145 cells expressing MEK5 shRNA (clone #7) (Fig. 2d) with similar results. Finally, ectopic expression of a MEK5 construct (Supplemental Fig. 1e) in PC3/shMEK5 (clone #12) cells showed that DNA-PKcs S2056 phosphorylation was restored to normal levels in response to irradiation, while phospho-ATM remained at similar levels between shMEK5 and shMEK5/MEK5 cells (Fig. 2e).

**Fig. 2:**
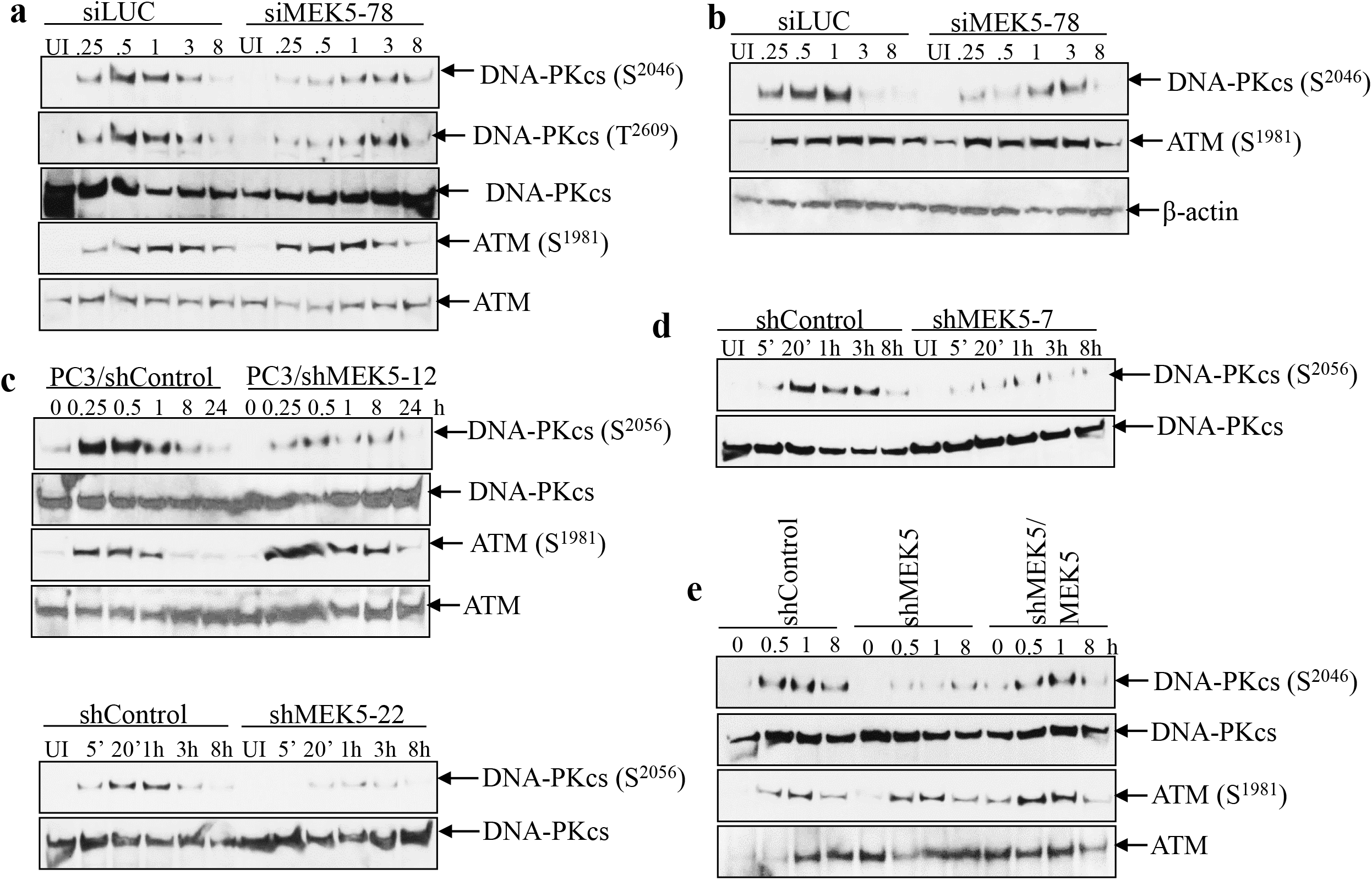
MEK5 knockdown impairs DNA-PKcs phosphorylation in response to ionizing radiation. PC3 (**a)** or DU145 (**b)** cells were transiently transfected with *Luciferase* siRNA (siLUC) or siRNAs against *MEK5* (#78). Four days later, cells were irradiated with 3 Gy γ-radiation, lysates were prepared at the indicated times and immunoblotted with the indicated antibodies. **c** PC3 cells stably expressing a control (shControl) or *MEK5* (clone#12, *upper*; clone#22, *lower*) shRNA were exposed to 3 Gy of γ-rays and cells were lysed at the indicated times. Lysates were immunoblotted sequentially with the indicated antibodies. **d** DU145 cells stable expressing a scrambled (shControl) or *MEK5* (clone#7) shRNA were exposed to 3 Gy of γ-rays and cells were lysed at different times and immunoblotted sequentially with anti-phospho-DNA-PKcs (S2056) and anti-total DNA-PKcs antibodies. **e** Ectopic expression of MEK5 restores activation of DNA-PKcs. PC3 cells stably expressing *shControl*, *shMEK5* (clone#12), or shMEK5 transiently expressing MEK5-pcDNA3 vector were exposed to 3 Gy of γ-rays and lysed at the indicated times. Lysates were immunoblotted sequentially with phospho-DNA-PKcs (Ser2056), total DNA-PKcs, phospho-ATM (Ser1981) and total ATM antibodies. UI: unirradiated.

We also examined the impact of MEK5 silencing on the response to IR of two additional cell lines, the non-tumorigenic prostate epithelial cells EP156T and the androgen-responsive LNCaP cells. In contrast to PC3 and DU145, MEK5 ablation did not have an impact on the phosphorylation of DNA-PKcs (S2056) and ATM (S1981) in response to IR (Supplemental Fig. 6a, b). Likewise, cell proliferation assay showed that MEK5 ablation did not sensitize LNCaP cells to IR (Supplemental Fig. 6c).

To validate the impact of MEK5 silencing on DNA-PKcs activation further, we exposed PC3 and DU145 cells to etoposide and phleomycin, two compounds that inflict cell damage by generating DNA double strand breaks, which are predominantly repaired by NHEJ [28]. We first performed a dose response study using various concentrations of etoposide with PC3/shControl and PC3/shMEK5-12 cells. As shown in Fig. 3a, PC3 cells with MEK5 knockdown were exquisitely sensitive to etoposide treatment compared with control PC3 cells. In a similar experiment, we treated PC3/shControl and PC3/shMEK5-12 cells with 10 µM etoposide for 16 h, removed the drug, and incubated cells for an additional 4 days after which we counted the cells. While total cell count of untreated PC3 expressing shMEK5 did not differ from control cells, etoposide-exposed PC3/shMEK5 showed an 80% reduction in cell numbers compared with etoposide-treated control cells (Fig. 3b). Furthermore, treating PC3 cells with 10 µM of etoposide resulted in a robust increase of DNA-PKcs phosphorylation at S2056 (Fig. 3c). In contrast, phospho-DNA-PKcs was significantly lower in PC3/shMEK5 cells for the whole time course (Fig 3c and Supplemental Fig. 7a). These results were further confirmed with DU145 cells, as well (Supplemental Fig. 7b). ATM activation was not different between control and *MEK5* knockdown PC3 cells. Finally, we treated PC3 and DU145 cells with 60 µg/mL of phleomycin for 2 h to generate DSBs, removed the drug, and incubated the cells in drug-free culture medium for up to 4 h. As seen with IR and etoposide, the expected increase in phospho-S2056 was observed only in the cells with normal levels of MEK5, but not in cells with MEK5 knockdown (Fig. 3d). Unlike DNA-PKcs, ATM activation in response to phleomycin was independent of MEK5 in both cell lines. Collectively, these results show that MEK5 is required for the activation of DNA-PKcs in response to DSB genotoxic stress, and thus MEK5 acts upstream of DNA-PKcs. However, ATM activation is independent of MEK5.

**Fig. 3:**
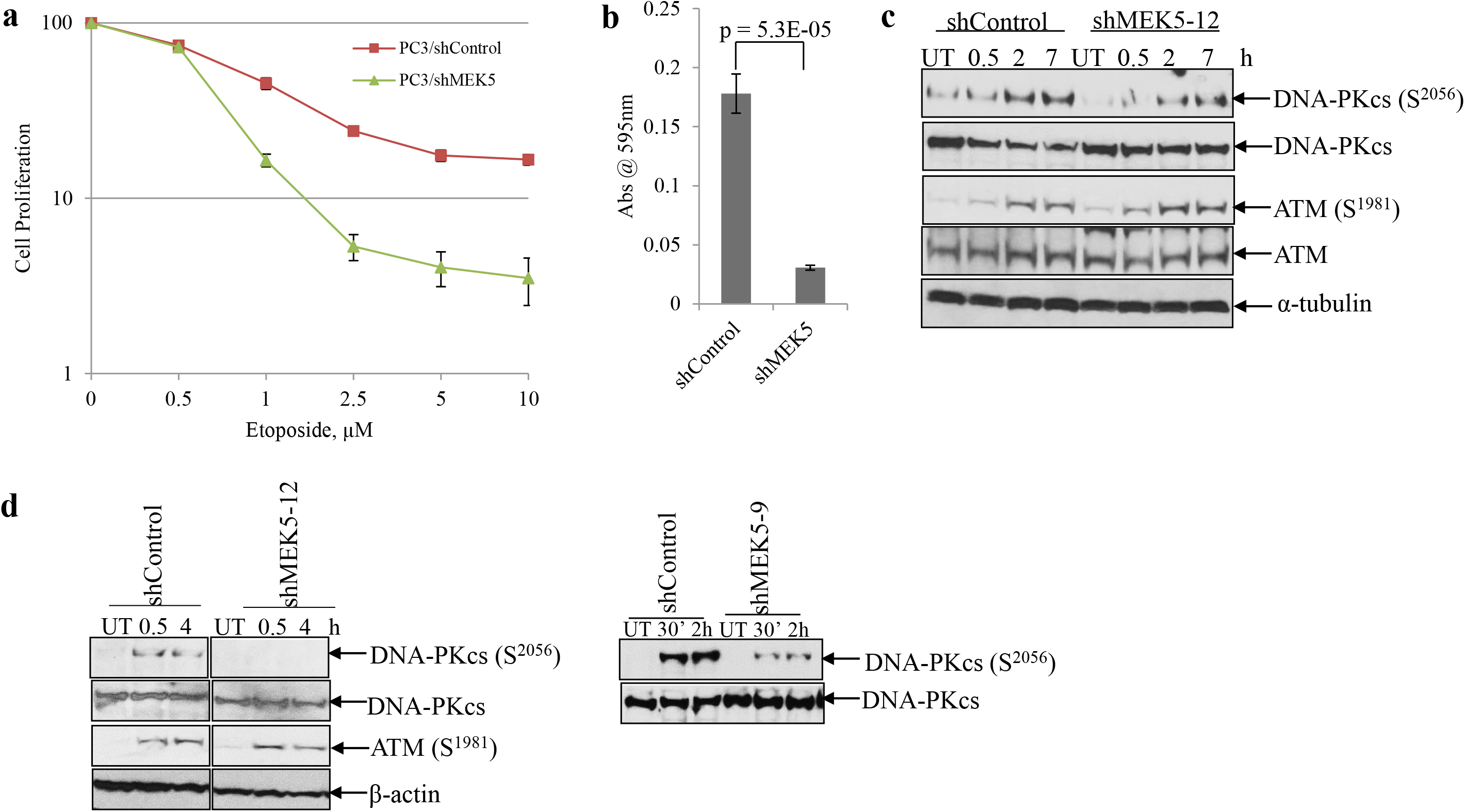
MEK5 knockdown impairs DNA-PKcs phosphorylation in response to etoposide and phleomycin. **a** dose response curves of control and *MEK5* shRNA in PC3 cells exposed to increasing concentrations of etoposide. Cell numbers were recorded 6 days post-treatment. **b** PC3 cells were exposed to 10 µM etoposide for 16 h, drug was removed, and cells were incubated for 6 days. Subsequently, cells were fixed, stained with crystal violet, stain dissolved and absorbance was measured at 595 nm. Mean ± S.D. (n = 3). **c** PC3 cells stably expressing a scrambled (shControl) or MEK5 (clone #12) shRNA were treated with 10 µM etoposide, cells were lysed at the indicated times and immunoblotted sequentially with the indicated antibodies. **d** PC3 cells (*left*) stably expressing *shControl* or *shMEK5* (clone #12) and DU145 cells (*right*) stably expressing *shControl* or *shMEK5* (clone #9) were treated with 60 µg/ml phleomycin for 2 h, drug was removed and cells were incubated for the indicated times. Lysates were immunoblotted with the indicated antibodies. UT: untreated.

### MEK5 ablation delays IR-induced foci resolution

An early response to DSBs is phosphorylation of H2AX, a variant of histone H2A, at serine 139, which is carried out by both ATM and DNA-PKcs [3]. Phosphorylated H2AX, called γH2AX, spreads from the double strand break over several megabases, and this can be visualized as foci by immunofluorescence using phospho-Ser139 antibodies. Similar to H2AX, 53BP1 is recruited to break sites and co-localizes with γHA2X. 53BP1 has been shown to be important for DNA repair by NHEJ [29]. To gain further insight into how MEK5 depletion sensitizes cells to genotoxic stress, we monitored the kinetics of γH2AX and 53BP1 foci formation in PC3 cells after exposure to 3 Gy γ-rays. The number of foci in unirradiated cells was low and it did not change with MEK5 silencing. As expected, radiation induced a rapid γH2AX and 53BP1 foci formation reaching maximum number within 30 min (Fig. 4a, c, d). MEK5 depletion (Fig. 4b) did not change the initial appearance of foci numbers. Subsequently, foci numbers in control PC3 cells were markedly diminished 2 h post-irradiation and returned to basal levels by 24 h. However, MEK5-depleted cells significantly delayed resolution of foci and they persisted above basal levels even after 48 h (Fig. 4a, c, d). We repeated the immunofluorescence experiments using transient *Luciferase* and *MEK5* siRNA transfection of PC3 cells with comparable results (Supplemental Fig. 8). Finally, we exposed PC3 cells to 10 µM etoposide and monitored H2AX and 53BP1 foci formation and resolution. In agreement with the IR treatment, exposure to etoposide resulted in increased number of foci at 30 min and 2 h, comparable for both control and MEK5 knockdown PC3 cells (Fig. 5a, b, c). However, foci resolution occurred much faster in control cells than in MEK5 silenced cells. We conclude that although the initial response to DNA damage is not dependent on MEK5 presence, the resolution and thus DNA repair of the damage is markedly delayed by MEK5 knockdown.

**Fig. 4.**
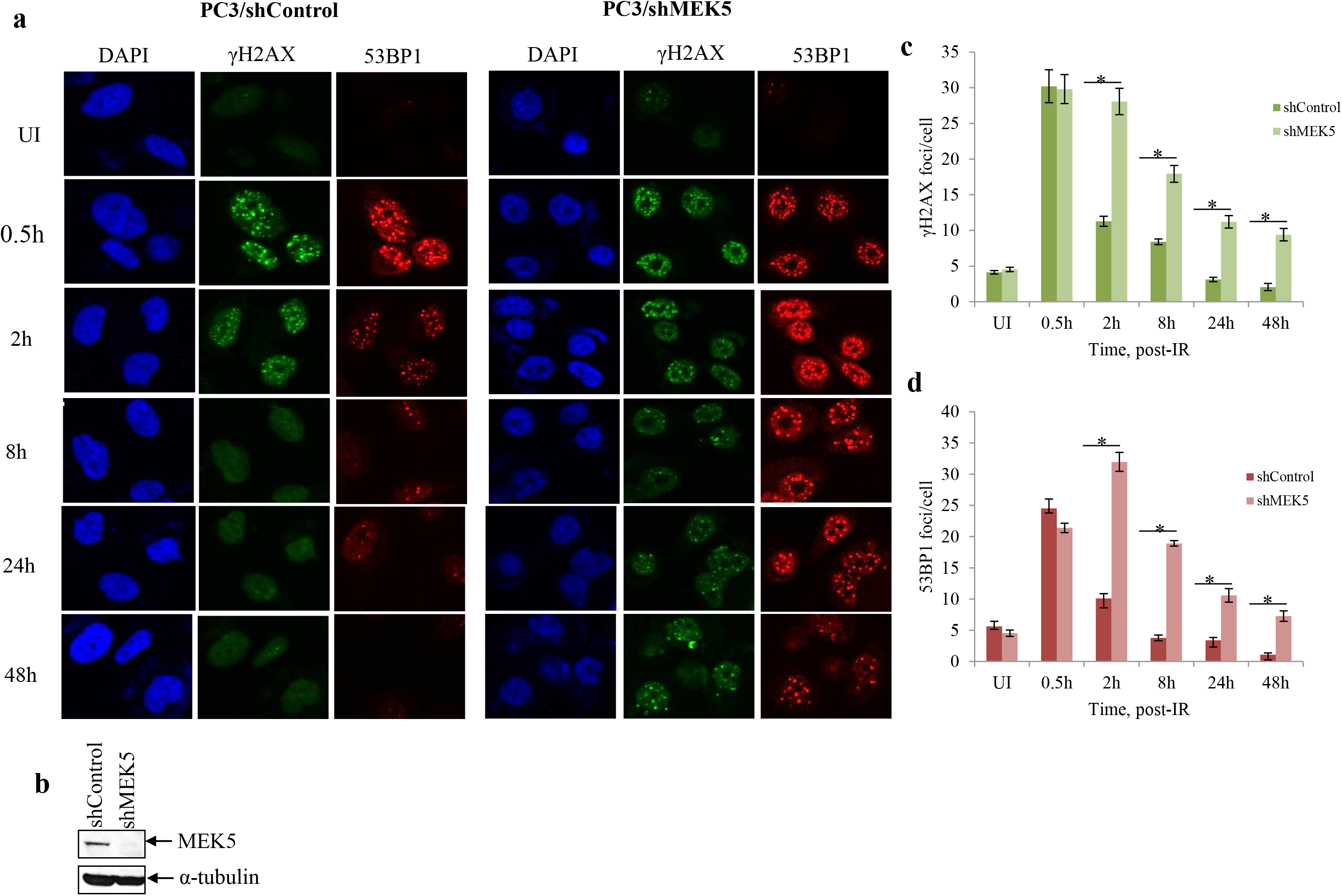
MEK5 knockdown delays resolution of irradiation-induced DSBs. PC3 cells stably expressing shControl or shMEK5 were exposed to 3 Gy γ-rays, fixed and stained for γH2AX, 53BP1, and 4’, 6-diamidino-2-phenylindole (DAPI; DNA). **a** Representative images and **b** western blot analysis of MEK5 protein levels in *shControl* and *shMEK5* (clone #12) cells. **c, d** quantitation of number of γH2AX (c) and 53BP1 (d) foci per cell over time after irradiation between cells expressing *shControl* and *shMEK5.* Shown mean ± S.D. (n = 3). * p < 0.001, calculated by Student’s t-test. UI: unirradiated.

**Fig. 5.**
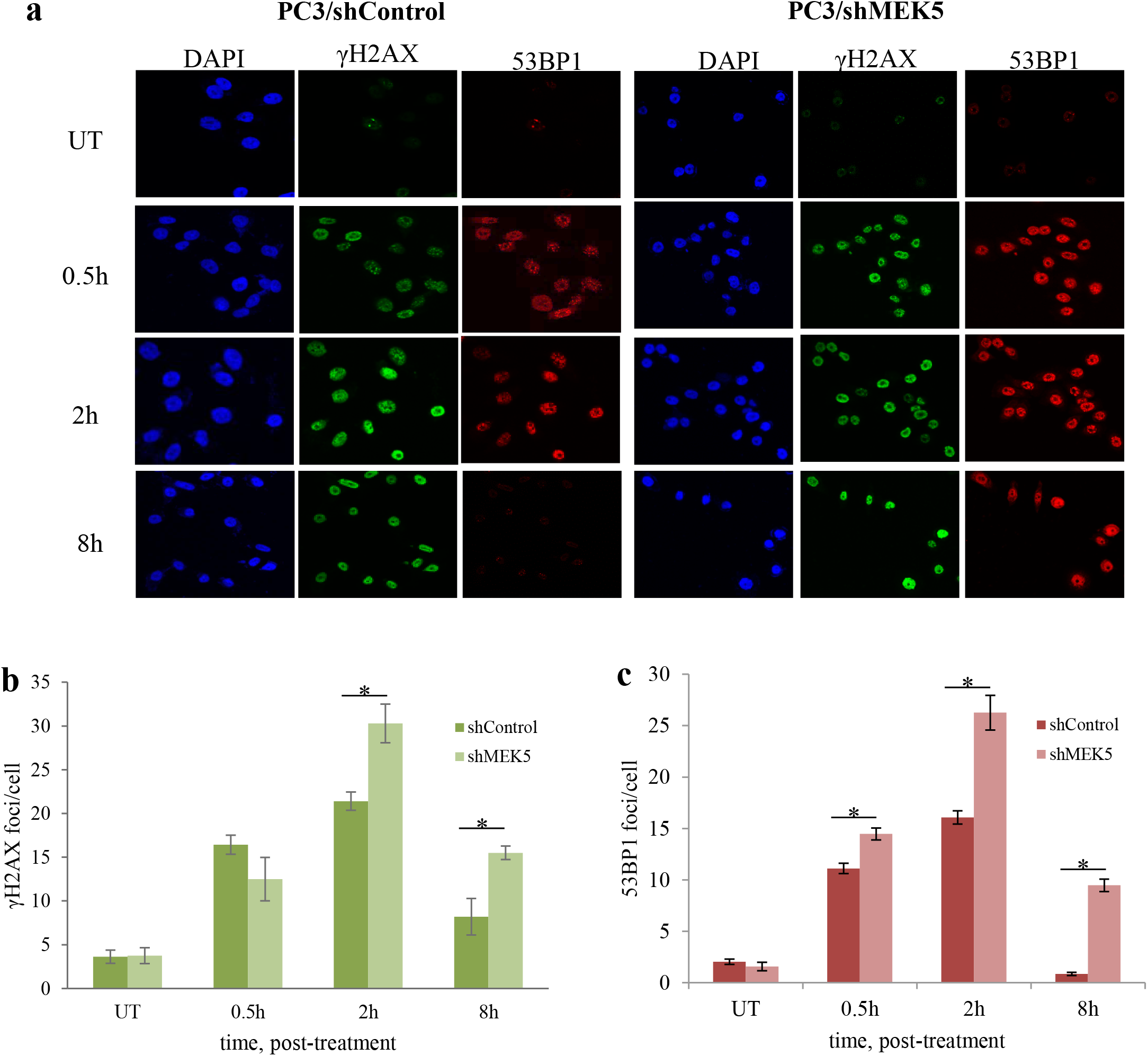
MEK5 knockdown delays resolution of etoposide-induced DSBs. PC3 cells stably expressing *shControl* or *shMEK5* were treated with etoposide and, at the indicated times, they were fixed and stained for γH2AX, 53BP1, and 4’, 6-diamidino-2-phenylindole (DAPI; DNA). **a** Representative images and **b, c** quantitation of number of γH2AX (b) and 53BP1 (c) foci per cell over time after irradiation between cells expressing *shControl* and *shMEK5*. Shown mean ± S.D. (n = 3). * p < 0.001, calculated by Student’s t-test. UT: untreated.

### MEK5 knockdown impairs non-homologous end joining

Next, we performed experiments to test directly the ability of MEK5 to promote NHEJ by using a cell-based assay [30, 31]. pEGFP-N1 plasmid (Clontech) was digested with HindIII restriction endonuclease (Fig. 6a) and transfected into PC3 cells expressing normal or reduced levels of MEK5. Transient transfection efficiency with the initial uncut plasmid was approximately 30% for both PC3 and PC3/shMEK5 cells as judged by the number of EGFP fluorescent cells measured under the microscope (Fig. 6b, *left*). Transiently transfected digested plasmid into PC3 cells resulted in approximately 10% of green fluorescent cells that express the protein. In contrast, PC3/shMEK5 produced almost 7 times fewer EGFP-producing cells (1.5%) (Fig. 6b, *right*). Thus, MEK5 downregulation reduces DNA-PKcs phosphorylation and impairs NHEJ.

**Fig. 6.**
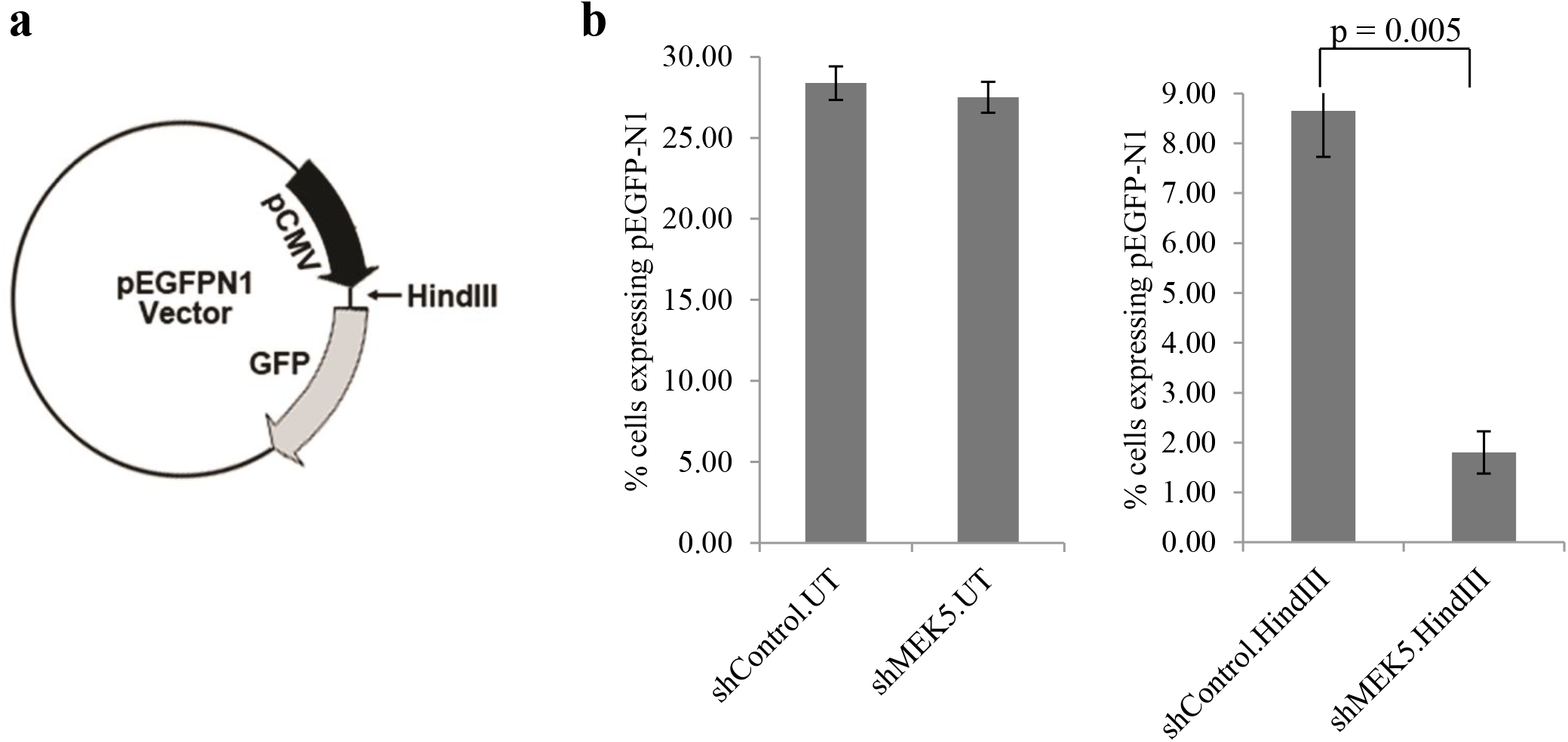
MEK5 depletion impairs non-homologous end joining. **a** Schematic of pEGFP-N1 vector with HindIII restriction site, **b** intact (*left*) or HindIII-digested (*right*) pEGFP-N1 vector was transiently transfected in PC3 cells expressing *shControl* or *shMEK5*. Twenty-four hours post-transfection, cells were fixed, stained with DAPI, and EGFP-positive cells were quantitated by fluorescence as percent EGFP-positive cells/total (DAPI) number cells. Mean ± S.D. (n = 3). P-value was calculated by Student’s t-test. UT: uncut plasmid; HindIII: restriction enzyme-digested plasmid.

### Combination of MEK5 blockade and ionizing radiation impairs tumor growth *in vivo*

To evaluate the efficacy of MEK5 knockdown combined with radiation to inhibit the growth of prostate cancer cells in mouse xenografts, we injected mice subcutaneously with PC3 cells expressing either shControl or shMEK5#12. We chose shMEK5 clone 12, as this clone showed the greater efficiency in downregulating endogenous MEK5 and, *in vitro* proliferation assays showed no appreciable difference in cell proliferation between shControl and shMEK5 PC3 cells (Supplemental Fig. 9). Mice bearing subcutaneous shControl or shMEK5 xenografts were either left untreated or exposed to a single dose of 4 Gy delivered specifically to the tumor by the Small Animal Radiation Research Platform (SARRP) irradiator using the onboard imager of the SARRP for image guided localization of the tumor (Supplemental Fig. 10) [32]. In agreement with *in vitro* proliferation assay, unirradiated shMEK5 cell growth showed a small but not significant (p = 0.5) impairment of growth when compared with unirradiated shControl cell growth. Likewise, exposure of shControl tumors to 4 Gy γ-rays had no effect on tumor growth compared with unirradiated shControl tumors (p = 0.5; Fig. 7). In contrast, shMEK5 cells exposed to radiation grew five-fold more slowly compared with unirradiated shMEK5 cells (p < 1E-04) (Fig. 7). In summary, these findings demonstrate that whereas MEK5 depletion or IR used separately have only a moderate impact on PC3 cells grown in mouse xenografts, the combination of MEK5 blockade with IR leads to a dramatic inhibition of tumor growth.

**Fig. 7.**
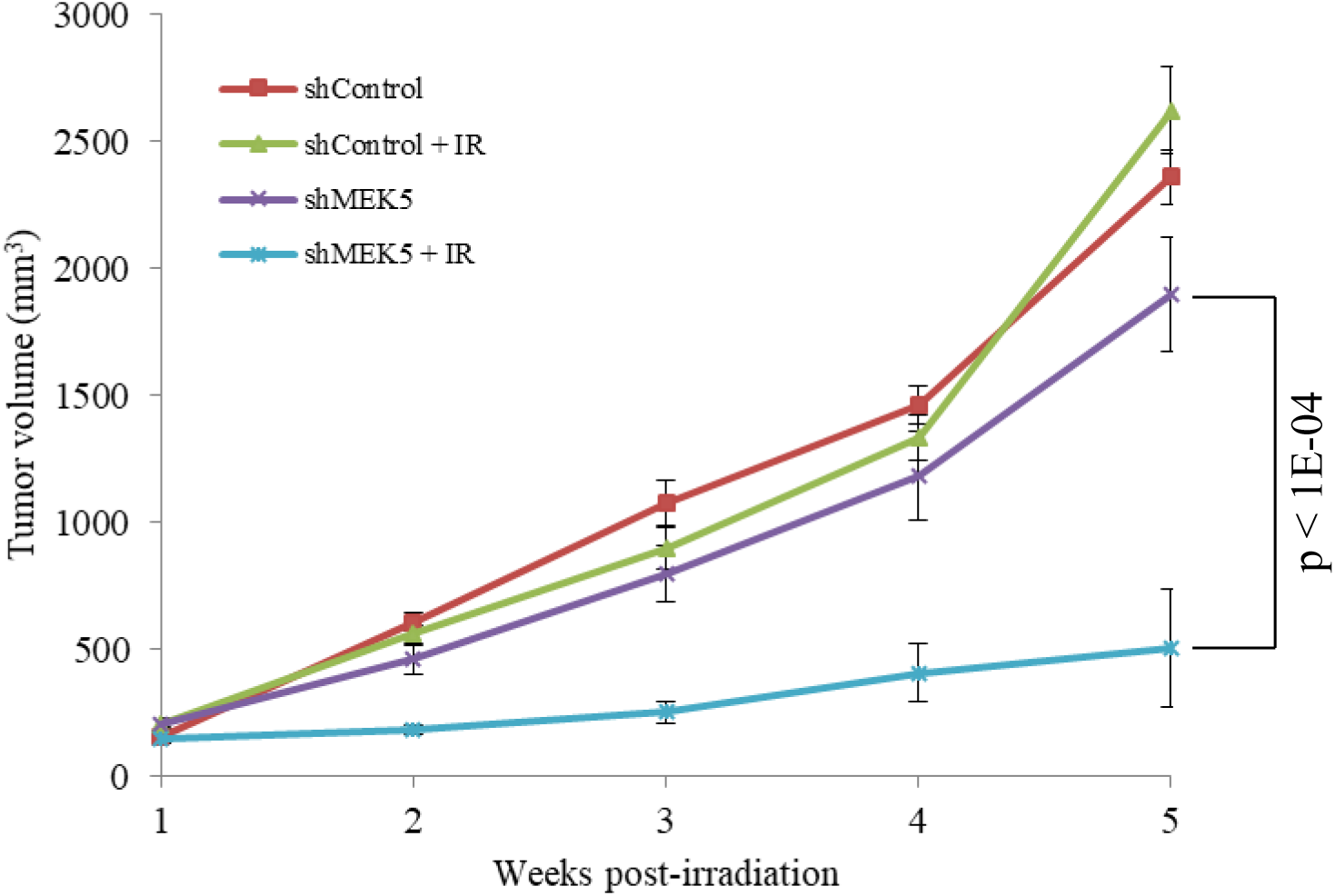
MEK5 ablation synergizes with radiotherapy to suppress PC3 tumor growth *in vivo*. PC3 cells stably expressing scrambled (control) or *MEK5* (clone #12) shRNA were injected subcutaneously into athymic male NU/J mice. When tumors reached ~200 mm^3^, mice were irradiated with 4 Gy x-rays (IR), or they were sham irradiated. Tumor growth was measured using a caliper. Shown mean volume ± S.E.M. (n = 8 mice /treatment).

## Discussion

In this study, we identified a critical role of MEK5 in mediating resistance to DNA damaging agents, such as ionizing radiation and etoposide, in prostate cancer cells. Our *in vitro* and *in vivo* studies demonstrate that MEK5 silencing sensitized PC3 and DU145 aggressive prostate cancer cell lines to IR, etoposide, and phleomycin through inactivation of DNA-PKcs and NHEJ repair.

In contrast, neither LNCaP nor EP156T cell lines were affected by MEK5 knockdown, most likely because these cells express relatively lower protein levels compared with PC3 and DU145. While ATM is activated by IR, etoposide, or phleomycin equally well between control and MEK5 knockdown cells, MEK5 silencing impairs phosphorylation of DNA-PKcs at S2056 and T2609 in response to genotoxic stress, indicating reduced activation. It has been shown that IR-induced DNA-PKcs autophosphorylation at S2056 is regulated in a cell cycle-dependent manner with attenuated phosphorylation in the S phase [8]. However, we confirmed that MEK5 silencing had no impact on cell cycle distribution and neither altered cell cycle arrest after IR. It is well established that elevated DNA-PKcs activity in various human cancers results in increased resistance to DNA damage. DNA-PKcs is associated with poor disease outcome [33] and predicts response to radiotherapy in advanced prostate cancer [34, 35], whereas knockdown of DNA-PKcs sensitizes DU145 and PC3 cells to ionizing radiation [36].

DSB generated by IR result in the formation of γH2AX and 53BP1 foci, and persistence of γH2AX foci indicate delayed repair and correlates with radiosensitivity [37–39]. The initial generation of IR-induced γH2AX and 53BP1 foci formation was similar between MEK5 knockdown and control cells. This can be attributed to ATM activation, which is known to play a dominant role in the generation of γH2AX, at least at early times post-irradiation [40]. In contrast, the resolution and thus repair of damage foci was markedly delayed in MEK5 knockdown cells compared with control cells. This is consistent with impaired DNA-PKcs action [40]. Furthermore, cell-based assays confirmed that NHEJ activity was significantly compromised in MEK5 knockdown cells. In the current study, we provide evidence for the first time that a member of the MAP kinase family, MEK5, has an impact on DNA-PKcs phosphorylation and NHEJ repair in response to genotoxic stress. Members of the mitogen-activated protein kinases (MAPK) family, especially the MEK1/2/ERK1/2 pathway, have been functionally associated with tumor DNA damage response and repair pathway, albeit with variable outcomes. Thus, activation of ATM by radiation downregulates phospho-ERK1/2, and this downregulation is associated with radioresistance in human squamous cell carcinoma cell lines [41]. Similarly, ERK1/2 activation in response to etoposide, which is abrogated in ATM knockout cells, leads to increased apoptosis and sensitization to the drug [42]. In contrast, ATM inhibition partly blocks phospho-ERK1/2 and diminishes HR in response to radiation, whereas inhibition of ERK1/2 activity reduced phosphorylation of ATM at S1981 in glioma cells [43]. Furthermore, treatment of pancreatic cancer cells with the MEK1/2-specific inhibitor trametinib resulted in significant radiosensitization by suppressing both HR and NHEJ [44]. In this case, it was noted that total DNA-PKcs levels were reduced in the trametinib-treated cells. However, inhibition of ERK1/2 has also been shown to increase DNA-PKcs activation and promote DSB repair by NHEJ in response to etoposide in breast cancer cells [45]. In our study, ERK1/2 activation in response to IR was not detected in PC3 cells (unpublished results). However, EGF treatment of PC3 cells was able to induce phospho-ERK1/2, implying that the MEK1/2/ERK1/2 pathway is intact in these cells. On the other hand, DU145 cells that constitutively express active ERK1/2 and phospho-ERK1/2 levels were not further induced by IR. Recently a study was published that showed ERK5 confers radioresistance to lung adenocarcinoma cell lines [46]. However, the mode of action of ERK5 in response to IR differs significantly from the present study. Thus, whereas ERK5 knockdown combined with radiation leads to compromised G2/M cell cycle arrest, our results show that MEK5 downregulation does not affect the cell cycle checkpoint response. Moreover, it was shown that IR caused sustained activation of ERK5, whereas we find that activation of ERK5 in prostate cancer cells is fast and transient reaching maximal levels of phosphorylation at around 10-30 min, diminishing thereafter and becoming undetectable by 2 hr post-irradiation. These differences may be attributed to different cancer types or, alternatively, to the fact that ERK5 has additional, MEK5-independent functions, and thus the impact of MEK5 knockdown may differ from that of ERK5 depletion [47].

In conclusion, our results support the mechanism that MEK5 inhibition sensitizes prostate cancer cells to genotoxic stress by severely impairing DNA-PKcs autophosphorylation and DNA repair by NHEJ. Our i*n vivo* experiments show that downregulation of MEK5 combined with irradiation markedly sensitizes prostate cancer cells to radiotherapy and support targeting MEK5 as a potential clinical intervention for intermediate and high-risk prostate cancer patients treated with radiotherapy.

## Materials and methods

Detailed experimental procedures describing cell culture, cell proliferation assays, irradiation, clonogenic survival assay, RNA interference and plasmid construction, cell cycle analysis, Western blot analysis, immunofluorescence, NHEJ assay, animal studies, and statistical analysis are included in the Supplementary Materials and Methods document.

## ACKNOWLEDGEMENTS

This work is supported by the Department of Defense Prostate Cancer Research Program, W81XWH-15-1-0296 (CGB). This study also used the resources of the Herbert Irving Comprehensive Cancer Center Flow Cytometry, Radiation Research, and Confocal and Specialized Microscopy Shared Resources funded in part through Center Grant P30CA013696. We thank Theresa Swayne and Laura Munteanu from the Confocal and Specialized Microscopy Shared Resource of the Irving Cancer Research Center (ICRC) and Drs. Siu-Hong Ho, Wei Wang, and Caisheng Lu from the Flow Cytometry Shared Resource of the Herbert Irving Comprehensive Cancer Center at Columbia University Irving Medical Center for their help in performing the immunofluorescence and flow cytometry experiments.

## Materials and Methods

### Cell Culture

Prostate cancer cells DU145, PC3, and LNCaP were grown at 37°C, 5% CO_2_ in RPMI 1640 (Invitrogen), supplemented with 10% fetal bovine serum (FBS; Thermofisher), 100 U/ml penicillin, 100 µg/ml streptomycin, and 2.5 µg/ml fungizone (Invitrogen). EP156T non-tumorigenic, immortalized, human prostate epithelial cells [1] were cultured in modified MCDB153 medium supplemented with 1% MEM non-essential amino acids solution, 200 nM hydrocortisone, 10 nM triiodothyronine, 5 mg/ml insulin, 5 mg/ml transferrin, 5 mg/ml sodium selenite, 100 ng/ml testosterone (Biological Industries USA), 5 ng/mL EGF, 50 mg/mL bovine pituitary extract, 100 U/ml penicillin, 100 mg/ml streptomycin and 1% fetal calf serum (Invitrogen) [2].

### Cell proliferation assay

Equal number of cells were seeded onto 6-well plates in triplicate. After treatment, cells were trypsinized and counted with a hemocytometer. Alternatively, cells were fixed in 4% paraformaldehyde for 20 min and stained with 0.1% crystal violet for 10 min. The stain was extracted by 10% acetic acid for 5 min and absorbance was measured at 595 nm using a spectrophotometer (Ultrospec 2000; Pharmacia Biotech).

### Irradiation

Subconfluent cell cultures were exposed to γ-rays at room temperature with the indicated doses by an Atomic Energy of Canada Gammacell 40 Cesium-137 Unit, providing a dose rate of 0.8 Gy/min.

### Clonogenic Survival Assay

To assess clonogenic survival, cells were transiently transfected with either *Luciferase* or *MEK5* siRNA. Forty-eight hours later, cells were trypsinized, counted, and added at 200 cells/well (DU145) or 400 cells/well (PC3) into 12-well plates in triplicate. Four to six hours later, cells were irradiated with 0, 2, 4, 6, or 8 Gy (DU145) or 0, 1, 2, 4, or 6 Gy (PC3) and incubated for 11 days. At the end of the incubation period, cells were fixed with 100% cold (−20°C) methanol for 20 min, washed once with PBS and stained with 0.5% crystal violet diluted in 20% methanol for 20 min. Colonies with more than 50 cells were counted under a microscope. The surviving fraction (SF) was calculated as (number of colonies formed)/ (number of cells seeded) X (plating efficiency).

### RNA Interference and Plasmid Construction

Stable depletion of MEK5 in PC3 and DU145 cells was achieved by infection with shRNA lentiviral particles (TRCN0000320675 MISSION shRNA Lentiviral Transduction particles; Sigma). Control shRNA expressing cells were established by infecting cells with scrambled shRNA lentiviral particles (SHC002V MISSION non-mammalian shRNA control transduction particles; Sigma). Down-regulation of MEK5 protein was assessed by Western blotting with MEK5 antibody (BD Biosciences). Selection of stable cell lines was performed with 1.25 µg/ml (PC3) or 0.5 µg/ml (DU145) puromycin for 2 weeks. Thereafter, cells were subcultured in medium containing the same concentration of puromycin.

RNA interference experiments were carried out with up to four non-overlapping siRNAs against *MEK5*, denoted #10, #20 [3], #76 (s11176; Ambion), and #78 (s11178; Ambion) and *Luciferase* (siLuc) that served as control [4], as described previously [4]. For transient transfection, cells at approximately 30% confluency were transfected with the indicated siRNA using Lipofectamine 3000 (Invitrogen), and protein abundance was examined 48–72 hr later by Western blot analysis. The human MEK5 construct was made by inserting *MEK5* cDNA, reverse-transcribed (SuperScript III, Thermofisher) from RNA isolated from PC3 cells, and cloned into BamHI/XhoI restriction sites of the pcDNA3 vector (Invitrogen). MEK5 was amplified using the following PCR primers: forward, 5’ – GGT ACT GGA TCC ATG TAC CCA TAC GAT GTT CCA GAT TAC GCT ATG CTG TGG CTA GCC CTT GGC CCC T – 3’ (contains a HA-tag) and reverse, 5’ -GGT AGA CTC GAG TCA CGG GGG CCC CTG CTG GCT CCG – 3’.

### Cell Cycle Analysis

Cells were harvested by trypsinization, washed in Dulbecco’s PBS (DPBS) and fixed in 70% ethanol overnight at −20°C. Fixed cells were subsequently washed in DPBS twice and incubated with propidium iodide/RNase staining buffer (BD Biosciences) for 30 min at 37°C. Flow cytometry was performed by a FACScalibur in conjunction with CellQuest software (BD Biosciences).

### Western Blot Analysis

Cells were washed with tris-buffered saline (TBS) and lysed in radioimmune precipitation assay (RIPA) buffer on ice for 30 minutes and then sonicated by 10 pulses at 30% amplitude using a Branson sonicator. Protein was subjected to either 3-8% Tris-glycine or 4–12% Bis/Tris (Invitrogen) SDS–PAGE. Quantitation of band intensity was performed with ImageJ software v1.47f (http://rsb.info.nih.gov/ij/). The following antibodies were used in this study: MEK5 antibodies were from BD Biosciences, DNA-PKcs, phospho-DNA-PKcs (S2056 and T2609) from Abcam, ATM, phospho-ATM (S1981), phospho-ERK5 (T218/Y220), ERK5 were from Cell Signaling, β-actin, and α-tubulin monoclonal antibodies from Sigma. γH2AX monoclonal antibodies were from Millipore, 53BP1 polyclonal antibodies were from Bethyl Laboratories.

### Immunofluorescence

PC3 cells were grown on 4-well chambers (EZ Millicel, Millipore). Cells were fixed with 4% paraformaldehyde in TBS for 20 min and permeabilized with 0.2% Triton X-100 in Tris-buffered saline (TBS) for 5 min. Nonspecific binding was blocked with 5% BSA in TBS containing 0.1% Tween-20 (TBST) for 1 h. Cells were double-stained with anti-γH2AX monoclonal antibody (1:500 dilution, overnight at 4 °C) and anti-53BP1 polyclonal antibody (1:1000 dilution, overnight at 4° C) followed by washing three times with TBST and incubation with anti-mouse secondary antibody conjugated to Alexa Fluor 488 and anti-rabbit secondary antibody conjugated to Alexa Fluor 594 (Thermofisher) (1:1000 dilution, 1 h, for both antibodies). Cells were washed once with TBST and incubated with 1 μg/ml 4′,6-diamidino-2-phenylindole (DAPI) for 7 min to stain the nuclei. Cells were washed three times with TBST and slides were mounted with ProLong antifade (Thermofisher) and stored at 4 °C protected from light. Images were acquired with a camera mounted on a Nikon Eclipse E600 fluorescence microscope. Foci numbers were calculated from at least 50 cells from 4 or more fields. Three independent experiments were performed.

### Non-homologous end-joining assay

A cell-based assay [5, 6] was used to measure NHEJ in PC3 cells expressing or not shMEK5. pEGFP-N1 plasmid (Clontech), which contains the enhanced GFP gene under the control of CMV promoter, was linearized with HindIII restriction endonuclease located between the promoter and the *GFP* gene coding sequencing. The HindIII digestion separates the CMV promoter from the coding sequence of GFP gene and thus it is unable to transcribe the gene. After digestion with HindIII, the linearized plasmid is separated and purified by agarose gel electrophoresis and Qiagen gel extraction kit (cat. no. 28704). Purified plasmid concentration was quantitated using Nanodrop One (Thermofisher). PC3 cells growing in 6-well plates were transiently transfected with 0.5 µg digested pEGFP-N1 plasmid. Eight hour later, cells were trypsinized and dispensed into 4-well microscopy chambers and allowed to attach and grow for an additional 16-18 h. Undigested plasmid was also transfected in parallel to ensure comparable transfection efficiency between PC3 and PC3/shMEK5 cells. At the end of the incubation, cells were fixed with 4% paraformaldehyde, permeabilized with 0.2% Triton X-100 in TBS and counterstained with DAPI, as outlined in the “Immunofluorescence” section of the Methods.

### Animal studies

All procedures were approved by the Columbia University Institutional Animal Care and Use Committee. Six-week old male athymic NU/J mice (Jackson Laboratory) were injected subcutaneously with 3×10^6^ PC3 cells expressing control (*shControl*) or MEK5 (*shMEK5*) shRNA mixed with growth factor-reduced matrigel (1:1). When tumors reached a volume of approximately ~200 mm^3^, mice were randomized to one of the following groups: (i) shControl, unirradiated; (ii) shControl; irradiated; (iii) shMEK5; unirradiated; (iv) shMEK5; irradiated. Mice were anesthetized with 100 mg/kg ketamide and 10 mg/kg xylazine in 0.9% saline by intraperitoneal injection and were either left untreated or irradiated with 4 Gy at a dose rate of 4 Gy/min using the Small Animal Radiation Research Platform (SARRP; Xstrahl, Suwanee GA) irradiator [7]. In particular, mice underwent cone beam computed tomography (CBCT) imaging using the onboard imager of the SARRP for image-guided localization of the tumor. A single beam was designed in the sagittal arrangement to deliver 4 Gy radiation through a 10×10 mm^2^ collimator prescribed to the isocenter. Radiation was delivered at a potential of 220 kVp and a filament current of 13 mA. Detailed radiation dosimetry and radiation planning information is provided in Supplemental Fig. 10. Tumor growth was measured with a digital caliper and the volume was estimated according to the formula *Length x Width^2^ x 0.50*, where length is the longest dimension and width the corresponding perpendicular dimension.

### Statistical Analysis

All in vitro experiments were carried out at least 3 times with similar results. Results from representative assays are shown. Data were represented as mean ± standard deviation (SD), unless otherwise indicated. P values were calculated by two-sided Student’s t-test, and values < 0.05 were considered statistically significant.

The dependence of tumor volume on treatment was modeled by analysis of variance (ANOVA) [8] as implemented in R [9, 10]. Outliers detected by the interquartile range test were discarded [11]. QQ-plots and the Shapiro-Wilk test showed that the samples were log-normally distributed [9]. Two-factor ANOVA showed a significant non-additivity between MEK5 and irradiation so that one-factor ANOVA was used [8, 9]. All against all comparisons were made, with multiple testing taken into account by Westfall’s method [12], as implemented in Multcomp [13].

---

**Supplemental Fig. 1.**
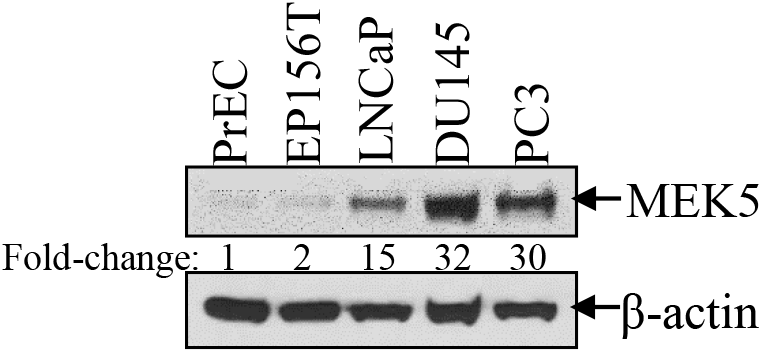
MEK5 protein expression in human prostate cancer cell lines. MEK5 protein abundance in a panel of normal (PrEC), immortalized (EP156T), and cancer (LNCaP, PC3, DU145) human prostate cell lines.

**Supplemental Fig. 2.**
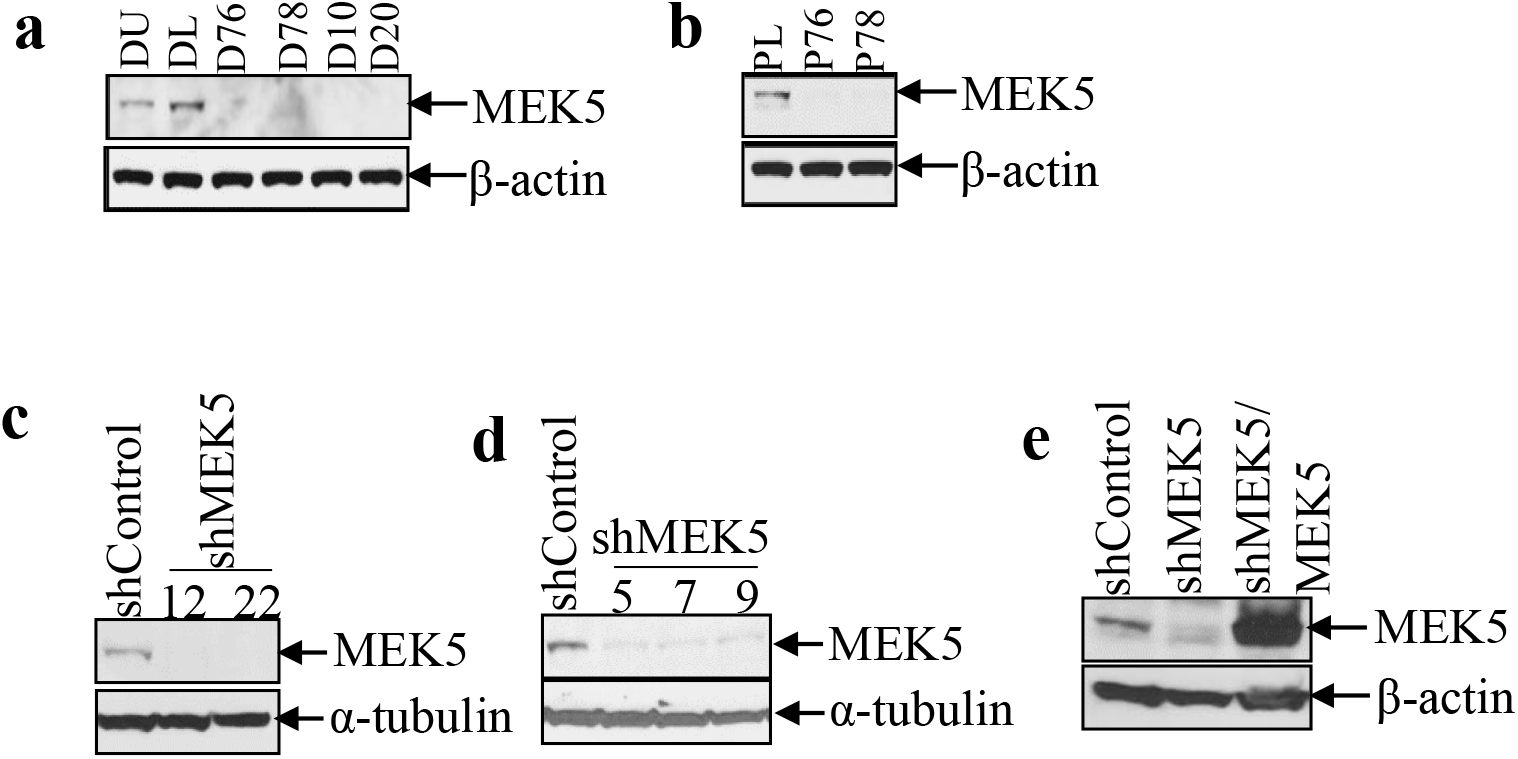
*MEK5* siRNA and shRNA efficiency. **a, b** MEK5 protein levels in DU145 (a) and PC3 (b) cells transiently transfected with *Luciferase* (DL or PL) or four (DU145; D76, D78, D10, D20), or two (PC3; P76, P78) independent *MEK5* siRNAs. **c, d** Stable clones of PC3 (#12, #22) (c) and DU145 (#5, #7, #9) (d) expressing *MEK5* shRNA or a scrambled (shControl) shRNA. **e** Ectopic expression of MEK5-pcDNA3 construct in PC3 stably expressing MEK5 shRNA (clone #12). Lysates were immunoblotted with anti-MEK5 and either anti-α-tubulin or anti-β-actin (loading controls).

**Supplemental Fig. 3.**
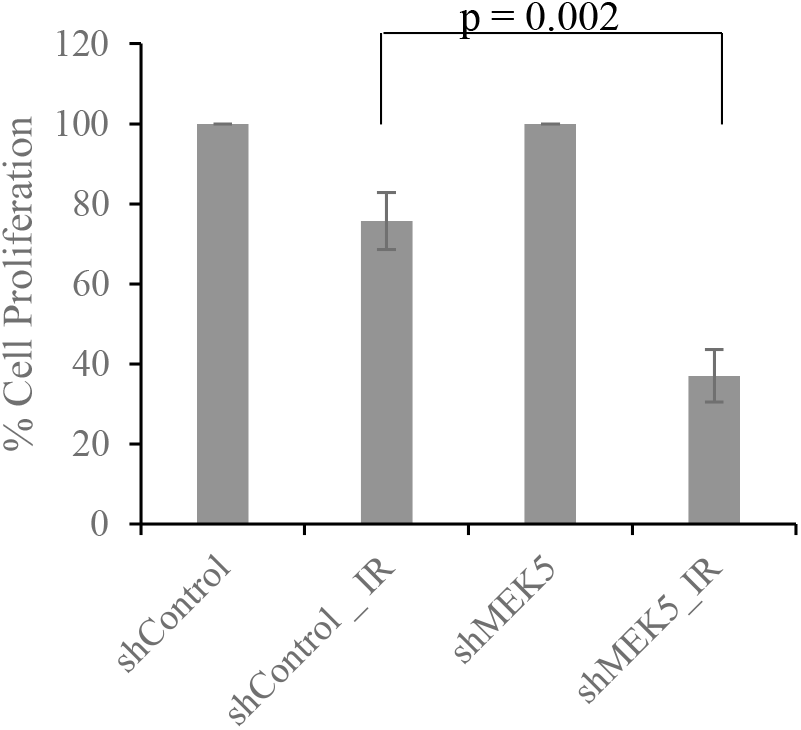
(Related to Fig. 1g) Equal number of DU145 cells stably expressing control or MEK5 (clone #9) were exposed to 4 Gy γ-rays or were sham irradiated. Cell proliferation was measured 5 days post-irradiation and expressed as percent proliferation relative to unirradiated cells. Shown mean ± S.D. (n = 3). P value was calculated by Student’s t-test.

**Supplemental Fig. 4.**
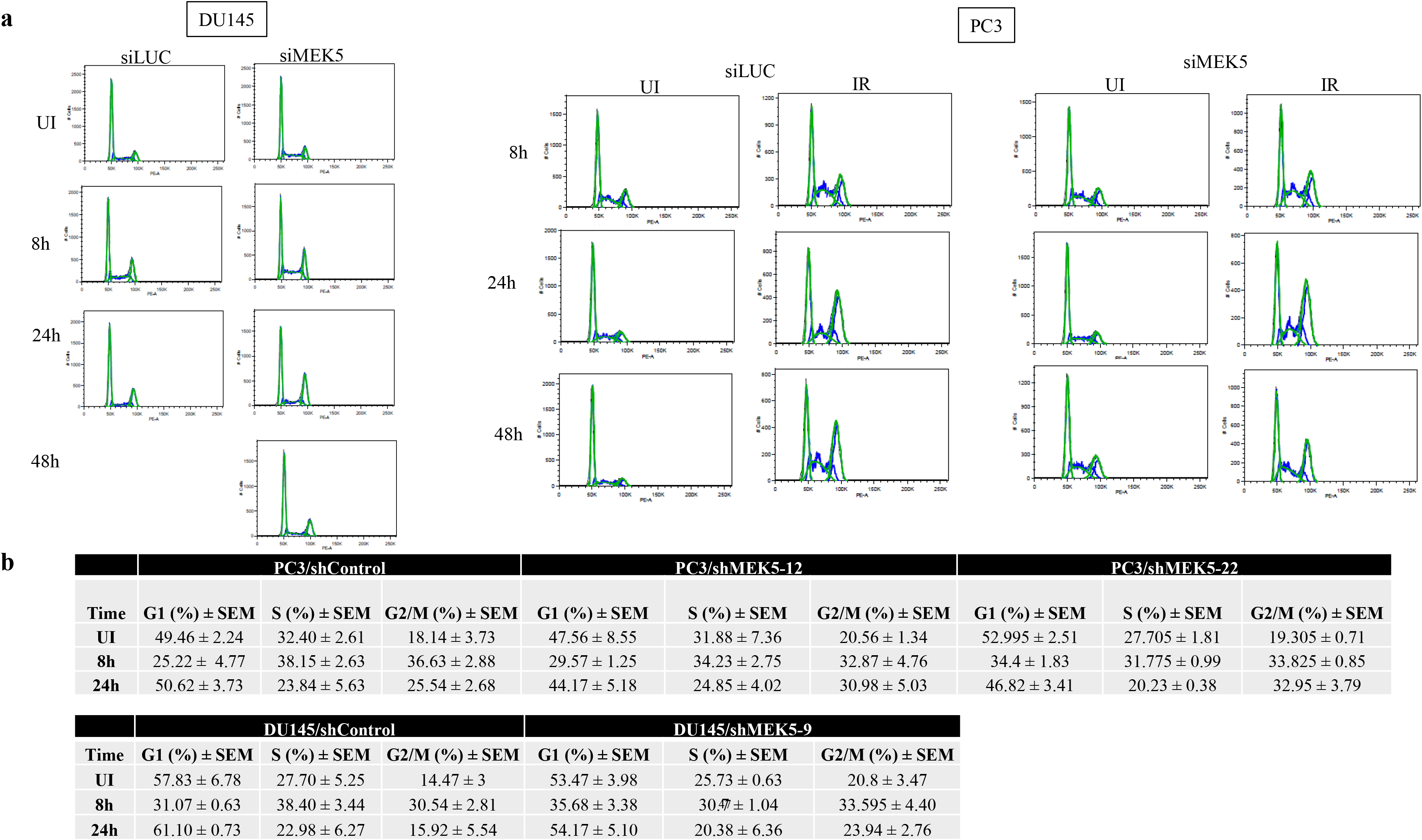
MEK5 downregulation does not influence cell cycle distribution within 48h of irradiation. **a** Cell cycle profile of DU145 (*left*) and PC3 (*right*) cells transiently expressing *Luciferase* or *MEK5* siRNA exposed to 3 Gy γ-rays or were sham irradiated (UI). **b** quantitation of cell cycle distribution of PC3 and DU145 cells stably expressing scrambled (shControl) or *MEK5* (PC3 clones #12 and #22; DU145 clone #9) shRNA. % G1, S, and G2/M phases were quantitated by FlowJo software. Shown mean ± S.E.M (n = 3).

**Supplemental Fig. 5.**
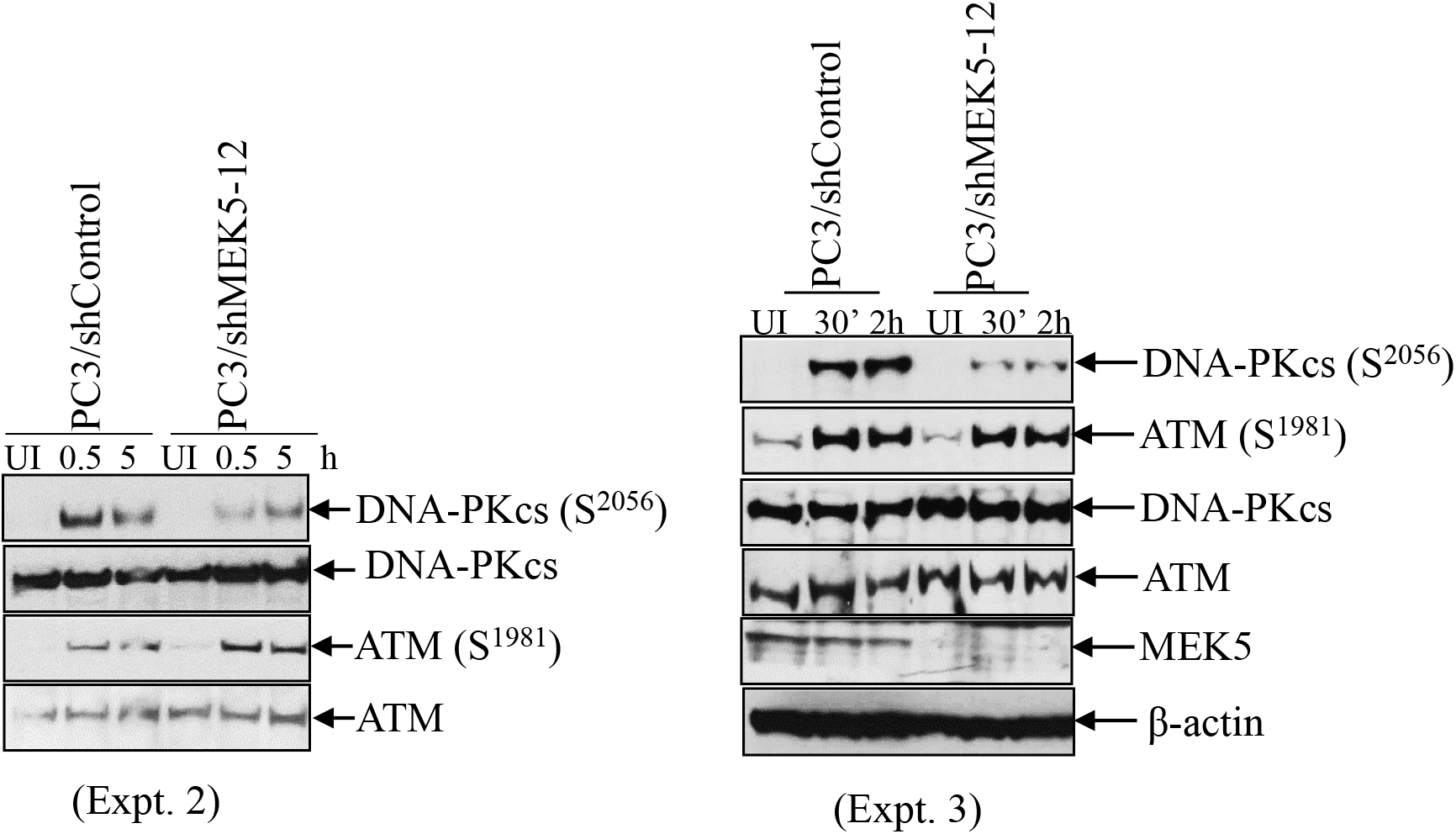
(Related to Fig. 2) Additional experiments (expt. 2 and expt. 3) with PC3 cells expressing *shControl* or *shMEK5* exposed to irradiation and immunoblotted for DNA-PKcs (S2056), ATM (S1981) phosphorylation. Blots were re-probed with antibodies against total DNA-PKcs, ATM, as well as MEK5 and β-actin (expt.3). UI: unirradiated.

**Supplemental Fig. 6.**
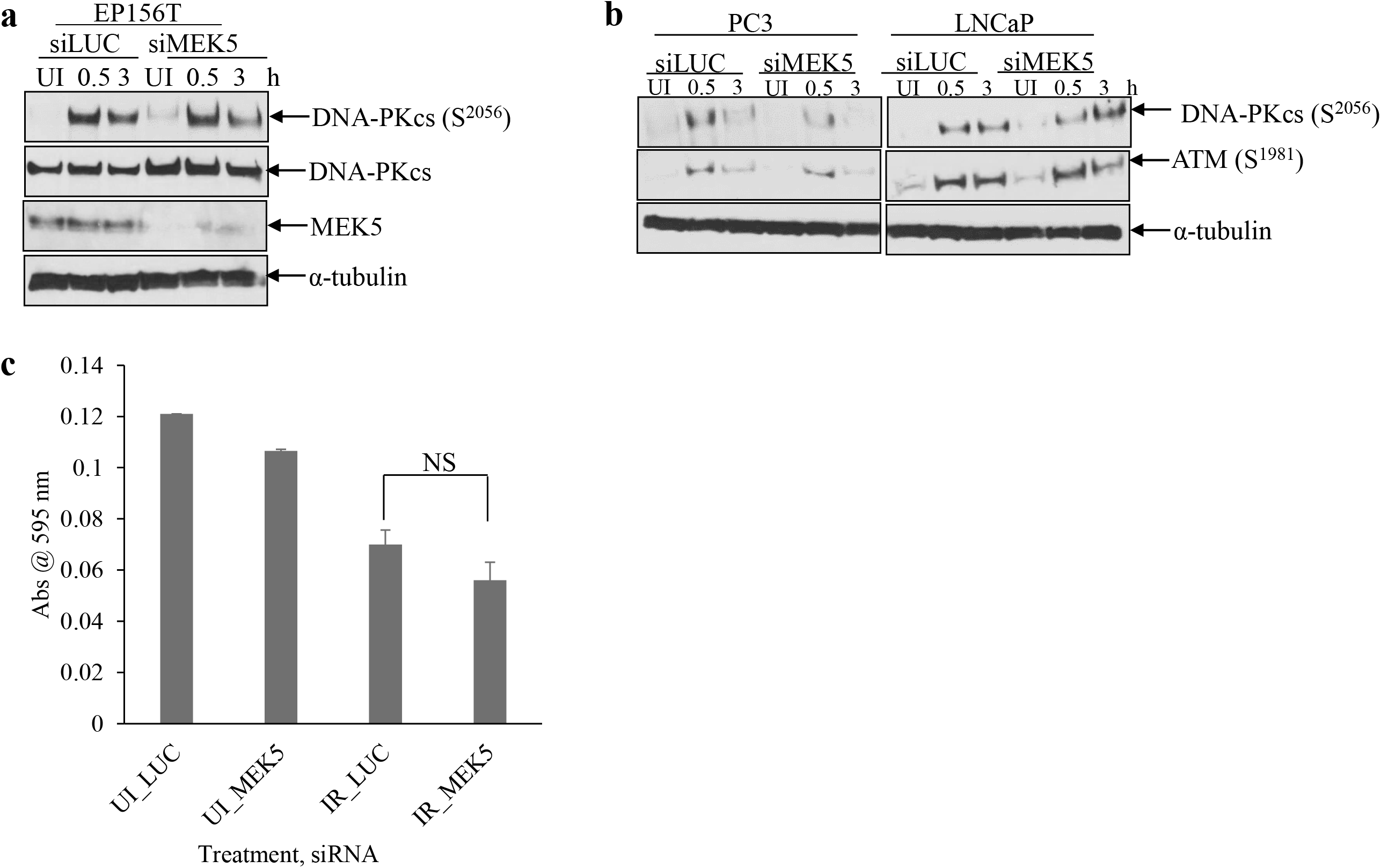
**a** EP156T cells or PC3 and LNCaP (**b**) were transiently transfected with *Luciferase* or *MEK5* (#78) siRNA and 6 days later they were exposed to 3 (PC3, EP156T) or 2 (LNCaP) Gy of γ-rays. Cells were lysed at various times and immunoblotted with the indicated antibodies. **c** LNCaP cell proliferation 6 days post-irradiation was measured by staining cells with crystal violet and recording absorbance at 595 nm. Mean ± S.D. (n =3) expressed as percent change compared with control unirradiated LNCaP cells. UI: unirradiated. IR: irradiated. NS: not significant.

**Supplemental Fig. 7.**
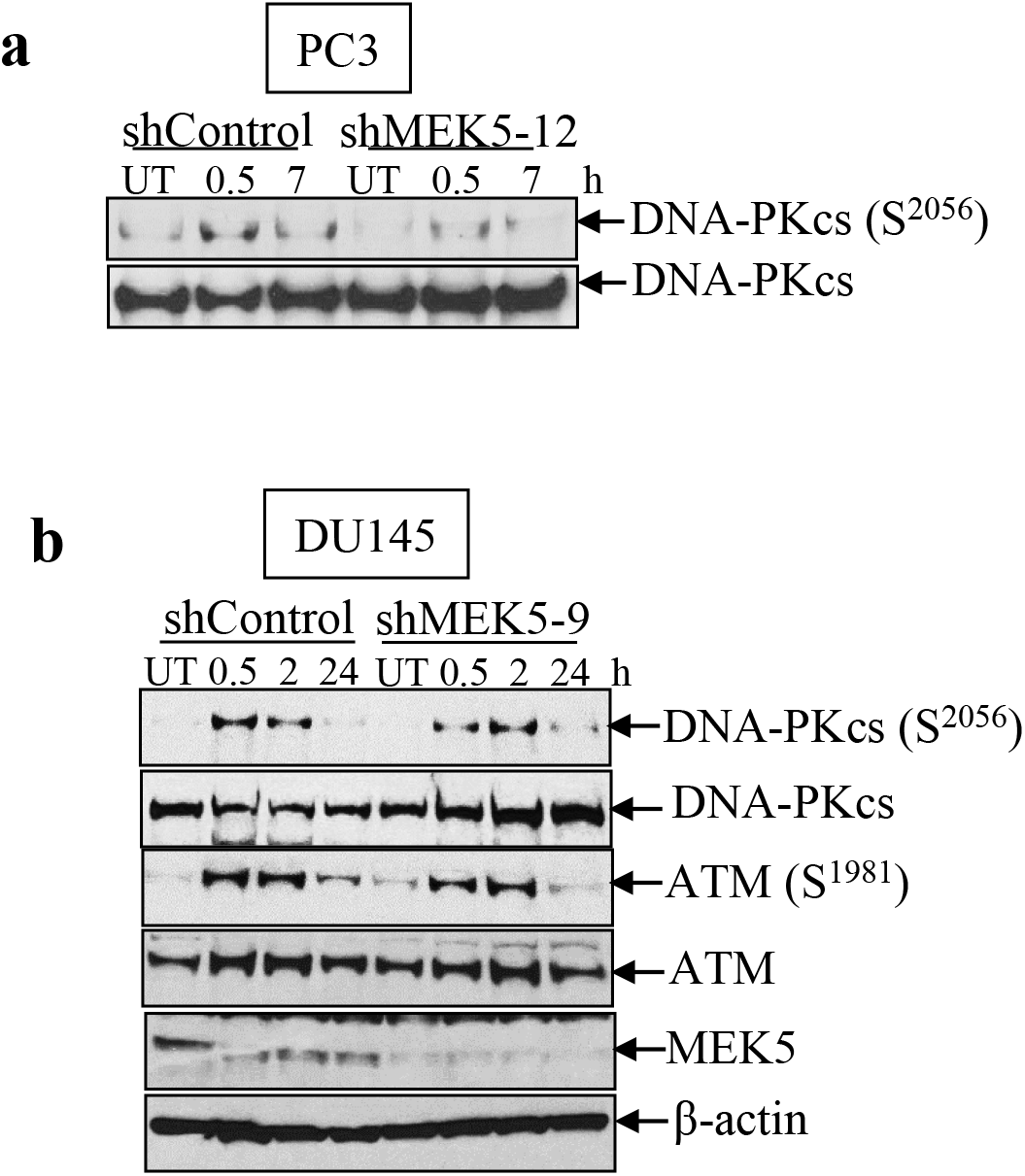
(Related to Fig. 3) PC3 (*shControl*, *shMEK5* #12) (a) or DU145 (*shControl*, *shMEK5*#9) (b) cells were serum-starved for 24h, then treated with etoposide (1 µM, 16h). The drug was removed, fresh medium (no serum) was added and cells were incubated for various times. Lysates were immunoblotted with the indicated antibodies. UT: untreated.

**Supplemental Fig. 8.**
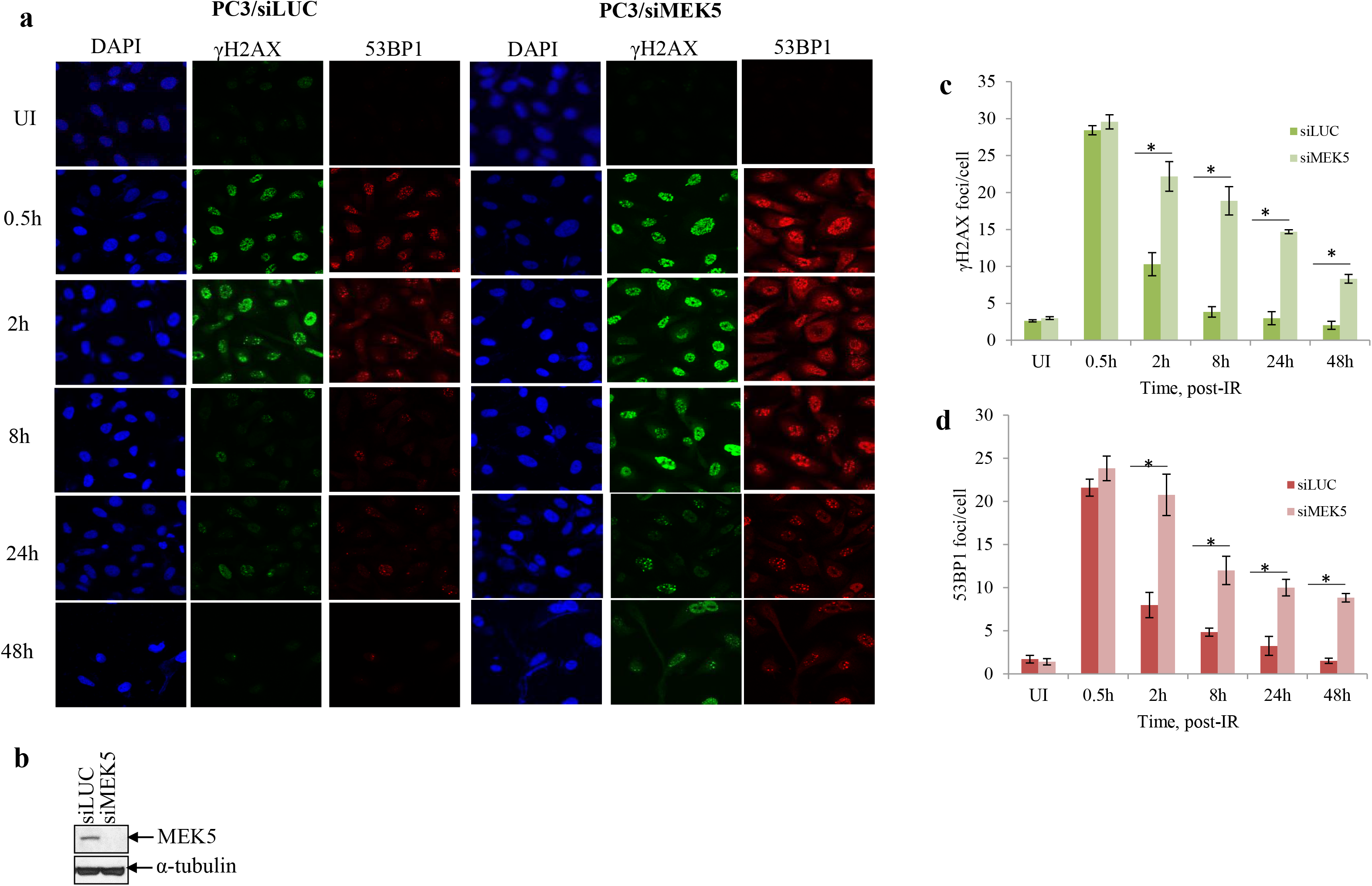
(Related to Fig. 4) PC3 cells transiently expressing Luciferase (siLUC) or MEK5 siRNA were irradiated with 3 Gy and at the indicated times they were fixed and stained for γH2AX, 53BP1, and 4’, 6-diamidino-2-phenylindole (DAPI; DNA). **a** Representative images and **b** western blot analysis of MEK5 protein levels in siLUC and siMEK5 cells. **c** quantitation of number of γH2AX foci per cell. **d** quantitation of number of 53BP1 foci per cell. Shown mean ± (n =3). * p < 0.001, calculated by Student’s t-test. UI: unirradiated.

**Supplemental Fig. 9.**
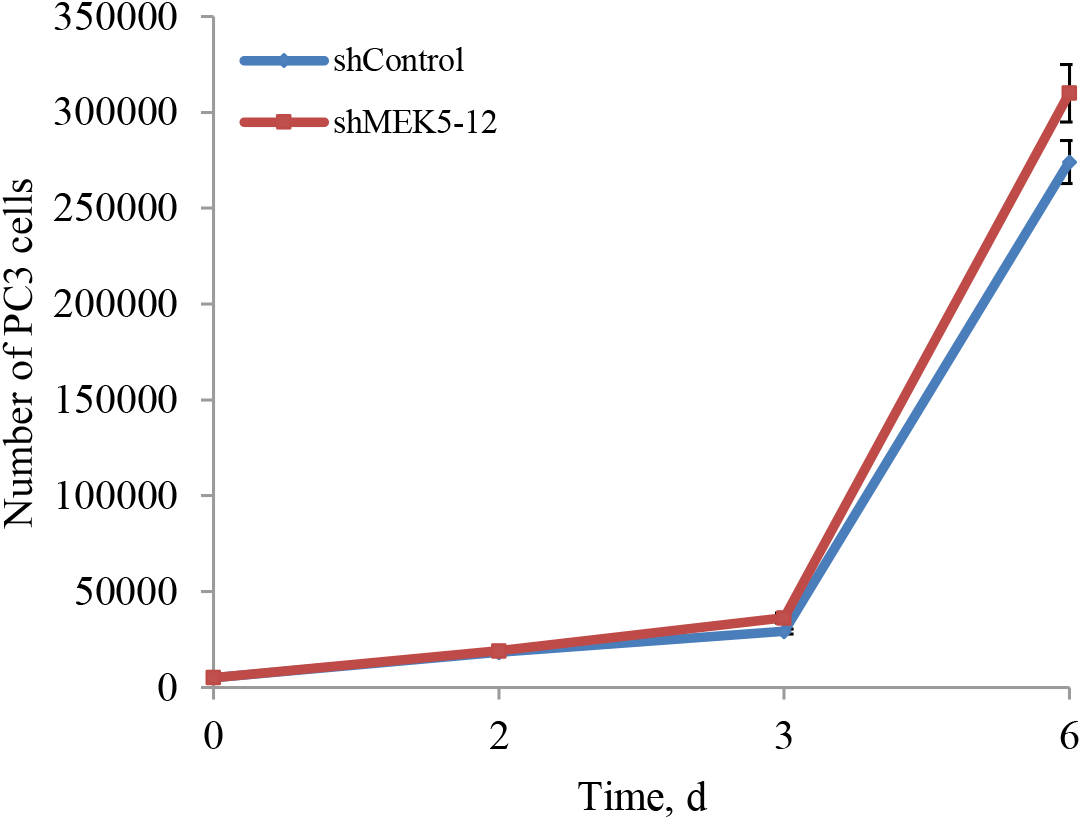
Cell proliferation. PC3 (shControl, shMEK5 clone#12) were seeded in 12-well plates (5,000 cells/well). Cells were trypsinized and counted with a hemocytometer at days 0, 2, 3, and 6. Mean ± S.D. (n = 3).

**Supplemental Fig. 10.**
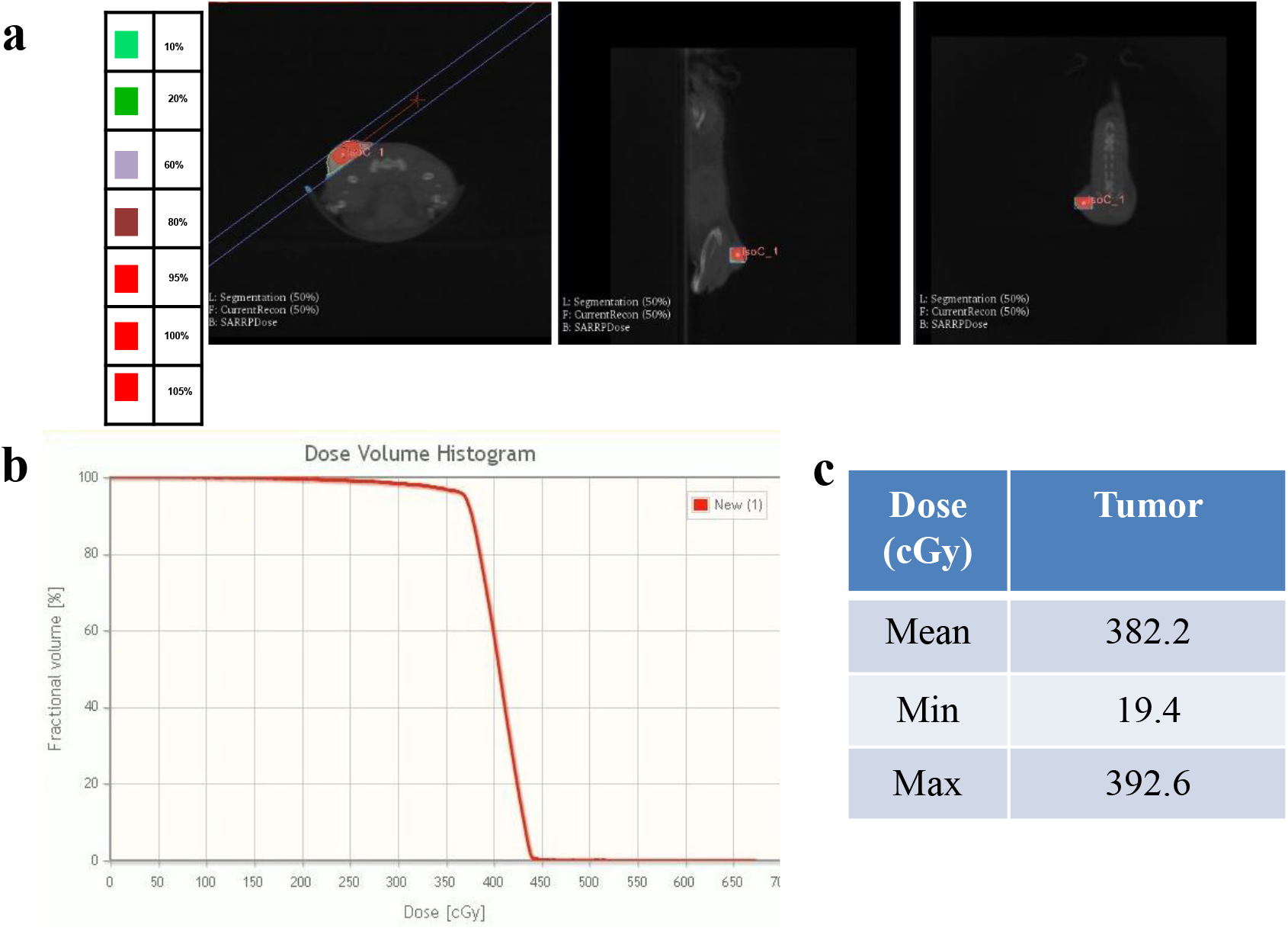
Radiation plan dosimetry and dose-volume histograms. **a** Representative images in coronal, axial and sagittal orientation of tumor-bearing mouse with radiation target volumes (tumor, red; tumor isocenter, cyan) contoured on cone-beam computed tomography images imported into MuriPlan software (Xstrahl). **b, c** Representative dose-volume histogram (DVH) and corresponding dosimetry to tumor (mean, minimum and maximum radiation dose in cGy) for tumor radiotherapy treatment plans.

